# Allele-specific activation, enzyme kinetics, and inhibitor sensitivities of EGFR exon 19 deletion mutations in lung cancer

**DOI:** 10.1101/2022.03.16.484661

**Authors:** Benjamin P. Brown, Yun-Kai Zhang, Soyeon Kim, Patrick Finneran, Yingjun Yan, Zhenfang Du, Jiyoon Kim, Abigail Leigh Hartzler, Michele L. LeNoue-Newton, Adam W. Smith, Jens Meiler, Christine M. Lovly

## Abstract

Oncogenic mutations within the epidermal growth factor receptor (EGFR) are found in 15-30% of all non-small cell lung carcinomas. The term exon 19 deletion (ex19del) is collectively used to refer to more than 20 distinct genomic alterations within exon 19 that comprise the most common EGFR mutation subtype in lung cancer. Despite this heterogeneity, clinical treatment decisions are made irrespective of which EGFR ex19del variant is present within the tumor, and there is a paucity of information regarding how individual ex19del variants influence protein structure and function. Herein, we identify allele-specific functional differences among ex19del variants attributable to recurring sequence and structure motifs. We build all-atom structural models of 60 ex19del variants identified in patients and combine ~400 μs of molecular dynamics simulations with biochemical and biophysical experiments to analyze three ex19del mutations (E746_A750, E746_S752>V, and L747_A750>P). We demonstrate that sequence variation in ex19del alters oncogenic cell growth, dimerization propensity, and tyrosine kinase inhibitor (TKI) sensitivity. We show that in contrast to E746_A750 and E746_S752>V, the L747_A750>P variant forms highly active ligand-independent dimers. E746_S752>V displays the least TKI sensitivity among the variants tested, which enzyme kinetic analysis and TKI inhibition experiments suggest is due to increased ATP Km relative to the common E746_A750 variant. Through these analyses, we propose an expanded framework for interpreting ex19del variants and new considerations for therapeutic intervention.

**Significance:** EGFR mutations are detected in approximately 30% of all lung adenocarcinomas, and the most common EGFR mutation occurring in ~50% of patients is termed “exon 19 deletion” (ex19del). Despite the existence of dozens of different genomic variants comprising what is generically referred to clinically as ex19del, clinicians currently do not distinguish between ex19del variants in considering treatment options, and the differences between ex19del variants are largely unstudied in the broader scientific community. Herein, we describe functional differences between distinct EGFR ex19del variants attributable to the structural features of each variant. These findings suggest a possible explanation for observed differences in patient outcomes stratified by ex19del subtype and reinforce the need for allele-specific considerations in clinical treatment decision making.

## Introduction

Oncogenic mutations within the epidermal growth factor receptor (EGFR) tyrosine kinase domain are detected in 15 – 30% of all cases of non-small-cell lung cancers (NSCLC) (1, 2). The two most common EGFR kinase domain (KD) mutations are a point mutation in exon 21, L858R, and a series of variants resulting in deletions within exon 19 (henceforward categorically referred to as ex19del mutations) (1, 2). More than a dozen genomic variants of ex19del have previously been identified (3, 4). Historically, ex19del mutations have not been differentiated in the clinic, and despite the known heterogeneity within this cohort of *EGFR*-mutant lung cancer, variant specific differences in ex19del have not been widely considered.

This is in stark contrast to the less frequently occurring EGFR exon 20 insertion (ex20ins) mutations. Several reports have described in detail the heterogeneity that different ex20ins variants display in terms of enzymatic activity and sensitivity to existing FDA-approved TKIs (5–9). At the structural level, molecular dynamics (MD) simulations suggest that ex20ins mutants can lower the free energy barrier associated with adopting the KD active conformation in an allele-specific manner (10). There are multiple ongoing drug development efforts aimed at designing TKIs to treat tumors harboring ex20ins variants in an allele-specific way (11–13). Preliminary assessment of an ongoing clinical trial (NCT03974022) suggests that this approach may be efficacious in ex20ins (14). The case has been made that we must evaluate drug efficacy on a per-mutant basis for ex20ins while ironically grouping all ex19del variants together (15). However, several retrospective studies have now suggested that there are differences in patient outcomes between ex19del patient populations (3, 4, 16–19). Emerging evidence suggests that structural classification of EGFR mutants can improve retrospective prediction of drug sensitivities (20). The lack of allele-specific resolution of ex19del variants in clinical practice may impede our ability to provide optimal therapeutic strategies for NSCLC and other cancer patients.

It is also noteworthy that investigations into ex19del often use the verbiage “exon 19 deletion” to refer to different allele variants, making it more challenging to functionally characterize them and develop appropriate therapeutic strategies. For example, the mechanism of activation of ex19del has been reported to be both ligand-independent (21–24) and ligand-dependent (25–27), and it is unclear to what extent the discrepancy is a result of the use of different experimental methodologies or different ex19del variants evaluated in previous studies. We have also previously found that the development of osimertinib resistance to the G724S mutant is dependent on the specific ex19del variant (28), suggesting that ex19del structural differences can have therapeutic implications. Thus, to maximize the efficacy of targeted therapies we need to refine our understanding of oncogenic variants at the atomic level.

In this study, we tested the hypothesis that sequence variation between EGFR oncogenic ex19del mutations can lead to allele-specific activation and TKI sensitivity. We probed the AACR GENIE database (29) and identified 60 unique ex19dels and built structural models of each variant. Next, we selected three of the most common variants predicted to be structurally distinct for detailed computational, biophysical, and biochemical evaluation: E746_A750, E746_S752>V, and L747_A750>P. Altogether, our results demonstrate that ex19dels are a functionally heterogeneous population with potentially unique considerations for optimal therapeutic targeting.

## Results

### Ex19del sequence variants cluster by chemical conservation and thus function

We first investigated the sequence heterogeneity of ex19del variants by probing the AACR GENIE database (29). We identified 60 variants and mapped these variants to the EGFR kinase domain (KD) (Figure 1; Table S1). Structurally, exon 19 corresponds to the ß3 sheet, ß3-αC loop, and N-terminal half of the αC helix (Figure 1A). All residues are numbered with respect to WT in the immature form (e.g., we reference L858R instead of L834R). We identified mutants ranging in size from a single residue deletion to a net eight residue deletion (Table S1). The starting and stopping points for the deletions predominantly occurred at residues E746, L747, A750, T751, S752, and P753, such that the length of the ß3-αC loop is the primary subject of sequence variation when compared to the ß3 or αC regions (Figure 1B; Table S1). The predominant mutations are E746_A750 (62.9%), L747_P753>S (7.4%), L747_T751 (5.2%), E746_S752>V (4.0 %), and L747_A750>P (3.7%) (Figure 1C; Table S1).

**Figure 1.**
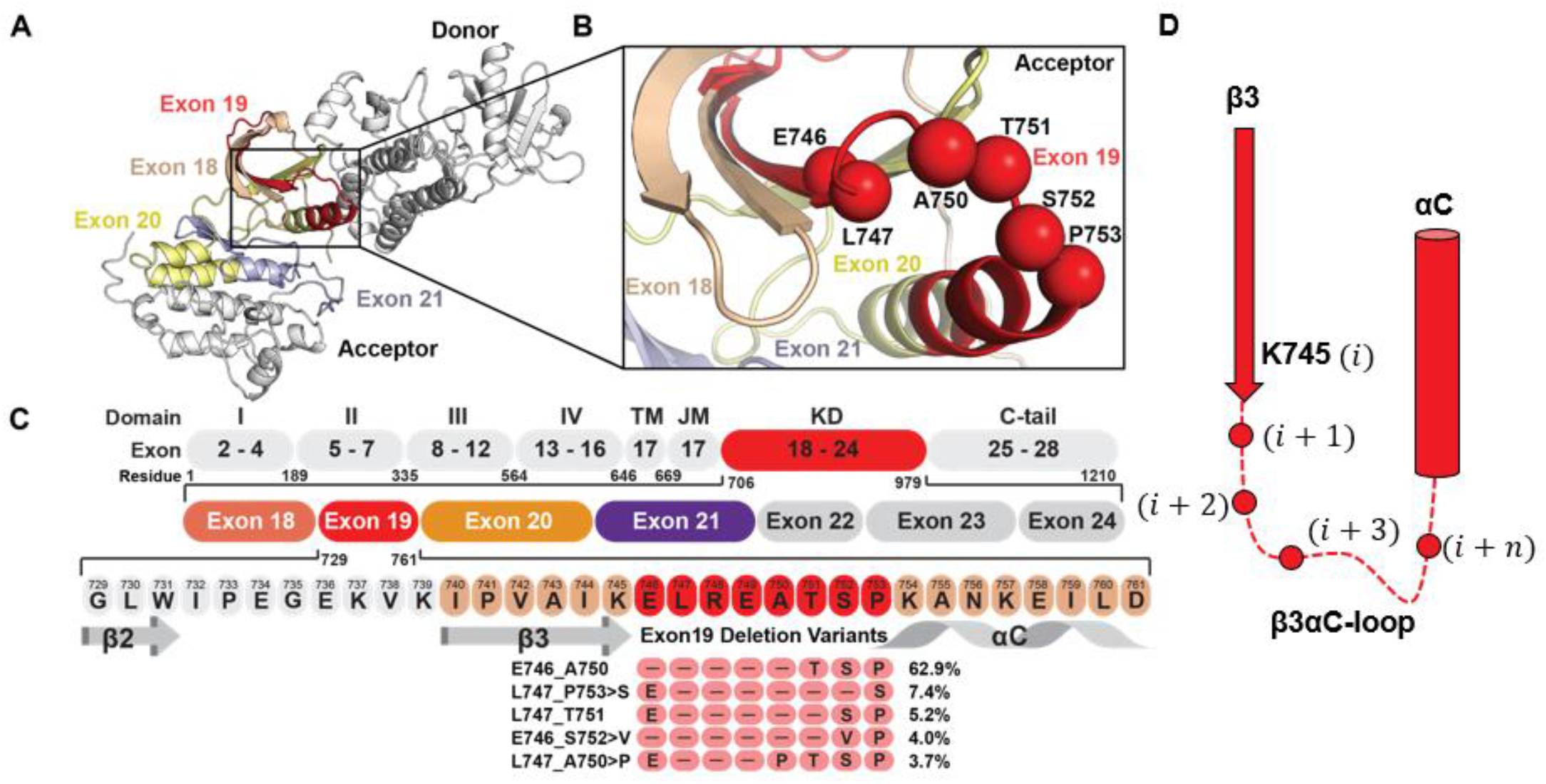
Frequently occurring mutations in the EGFR ß3-αC motif. **(A)** Schematic representation of the active EGFR-WT asymmetric dimer. Oncogenic and TKI resistance mutations have been reported in exons 18 (wheat), 19 (red), 20 (yellow), and 21 (blue). **(B)** The majority of deletion mutations begin at residues E746, L747, or T751. Deletion mutants frequently terminate with or without an insertion at position A750, T751, S752, or P753. Spheres indicate the residue Cα. **(C)** Multiple sequence alignment of the ß3-αC motif between EGFR-WT and ex19del variants with >2% frequency. **(D)** Residues at the β3αC interface can be referenced with respect to their index after the conserved K745 residue in the majority of mutants.

The breadth of variants is substantial, ranging from deletions that occur entirely in β3 (K739_I744>N) to those occurring almost entirely in αC (e.g., P753_I759). To help characterize the mutations, we first built structural models of all variants utilizing the Rosetta comparative modeling approach coupled with Gaussian accelerated MD (GaMD) (30) (see **Methods**). Our models suggested several recurring structural features of ex19del. First, the most common ex19del variants, including E746_A750, L747_P753>S, and L747_T751 (Figure 1C), replace L747 at the β3-αC interface with a serine and simultaneously remove at least one full turn from the N-terminus of the αC helix (Figure S1A). Second, mutants with net deletions of size three, such as L747_A750>P and E746_T751>APS, frequently converge on the same β3-αC loop conformation, characterized by a ß3-αC tight turn with proline in the second position (Figure S1B). Third, we observed that several mutants project polar residues into the ATP binding pocket in the vicinity of the canonical K745 – E754 salt bridge, such as L747_S752>Q and E746_S752>V (cis-trans proline-dependent).

To evaluate potential functional differences between mutants, we selected three isoforms that are prevalent in patients based on our AACR GENIE analysis (Figure 1C) and cover the breadth of features described above: E746_A750 (62.9%), E746_S752>V (4.0%), and L747_A750>P (3.7%). For clarity, we periodically reference residues by their position relative to K745 (Figure 1D).

### Ex19del variants adopt unique ß3-αC conformations with different energetic barriers to activation

We began with the hypothesis that ex19dels can display allele-specific differences in their propensity to adopt the active conformation. Wild-type EGFR (WT) is activated when ligand binds the extracellular domain (ECD) to promote intermolecular dimerization and further oligomerization (31–33). Intracellularly, these conformational changes result in asymmetric dimerization between two KD where the “receiver” KD is stabilized in an active conformation by the “donor” KD (34). Previous investigations have shown that oncogenic variants in the KD often stabilize the αC-helix by suppressing intrinsic disorder (35) leading to enhanced dimerization where the mutant KD behaves as a “super acceptor” (36).

Subsequently, we performed six independent conventional molecular dynamics (cMD) simulations of 4.0 – 6.0 μs for each mutant and state (E746_A750, E746_S752>V, and L747_A750>P, in active and inactive state respectively), such that three simulations were initiated from each state (120.0 μs total). Consistent with previous reports (37), the αC helix of WT readily departed from the active conformation to adopt an unstructured intermediate state, and 1/3 active state simulations transitioned completely to the Src-like inactive conformation (αC helix out, A-loop in, DFG in) (Figure S2A; Movie S1).

In comparison, each of the ex19del variants were stabilized in the active state (αC helix in, A-loop out, DFG in, Figure S2B – D). The tight turn predicted in the Rosetta/GaMD model of L747_A750>P is restricted in its motion, preventing inactivation (Figure S2D). Unfortunately, no transitions were observed from the inactive to the active state or *vice versa* in any of the ex19del cMD simulations. Therefore, we combined steered MD (SMD) with umbrella sampling (UMD) simulations to map the conformational free energy landscape (FEL) of the transition.

Following a procedure similar to that previously employed for ex20ins variants (10), we defined our umbrella sampling collective variables (CV) along two dimensions: (1) Activation state of the αC helix as defined by the difference in distance between K860 – E762 and K745 – E762, and (2) activation state of the A-loop as defined by the dihedral angle formed by the Cα atoms of D855 – F856 – G857 – L858 (Figure 2A, B).

**Figure 2.**
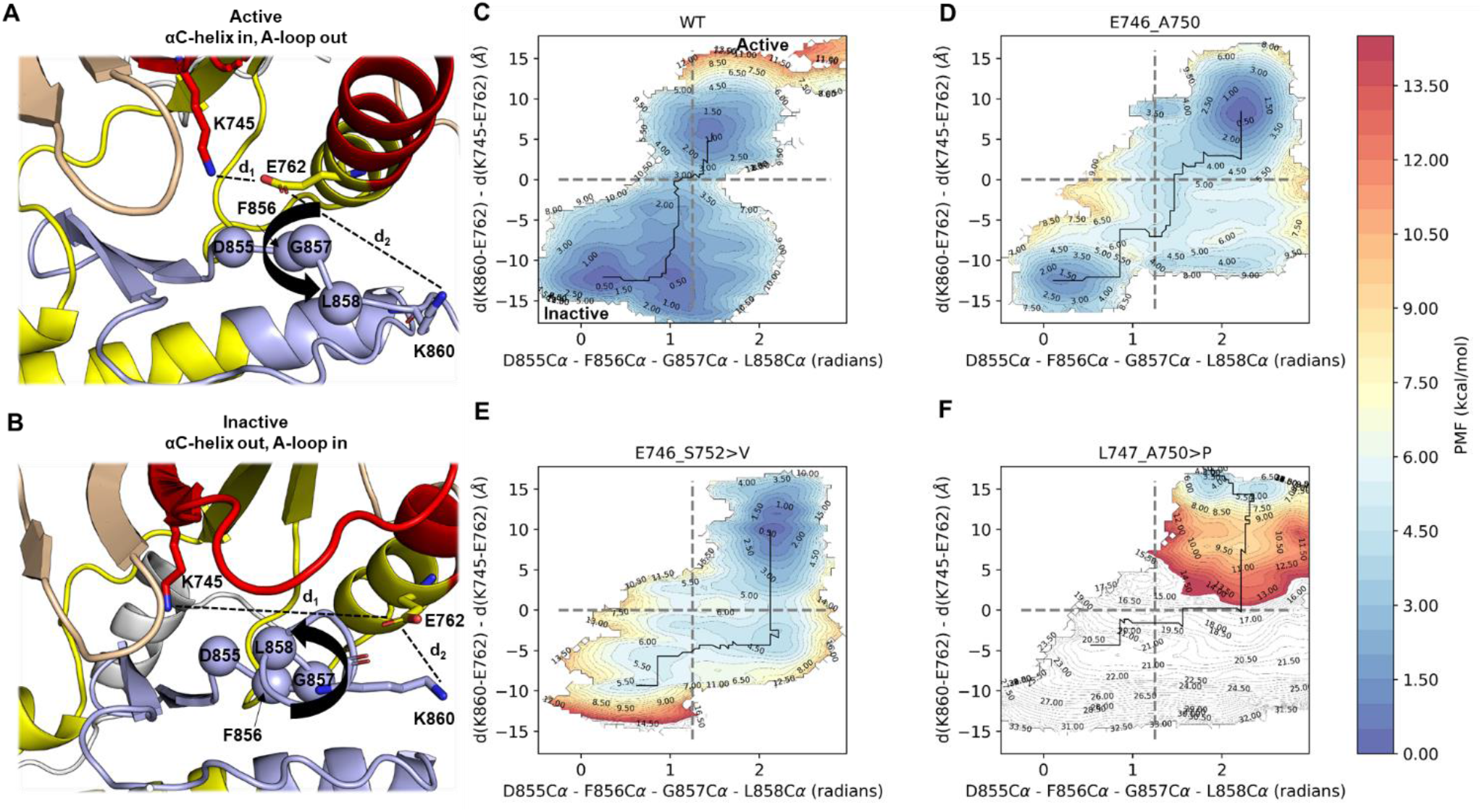
Conformational free energy landscapes of ex19del variants from umbrella sampling MD simulations. Collective variables describe the **(A)** active and **(B)** inactive states as the pseudo-dihedral angle formed by the alpha carbon atoms of residues D855, F856, G857, and L858 (x-axis) as well as the difference in distance between the capping sidechain atoms of E762 and K745 (d1) and E762 and K860 (d2) (y-axis). Conformational free energies are shown for **(C)** WT, **(D)** E746_A750, **(E)** E746_S753>V, and **(F)** L747_A750>P. Plots are contoured at 0.5 kcal/mol and colored within the range 0 (blue) and 15 (red) kcal/mol. Contours above 15 kcal/mol are colored white.

Using these 2 CVs, we measured the free energy difference between the active and inactive states of WT and found it to be approximately 1.0 kcal/mol in favor of the inactive state (Figure 2C), in good agreement with prior estimates (10). In contrast to WT and the previously reported exon 20 insertion mutations (10), all three ex19del variants favored the active state (Figure 2D – F). E746_A750 and E746_S752>V favored the active state by approximately 1.0 kcal/mol and 4.5 kcal/mol, respectively (Figure 2D – E). We also performed SMD+UMD simulations on the other two most commonly occurring ex19dels, L747_P753>S and L747_T751. L747_T751 displays an activation profile similar to E746_S752>V, while L747_P753>S may be more comparable to several ex20ins variants (10) (Figure S3).

Interestingly, L747_A750>P appears to be trapped in the active state, with prohibitively large free energy barriers to the inactive state (Figure 2F). We considered that this may be a result of the proline substitution at position 747. We tested this hypothesis by building models for the oncogenic missense variant L747P (38) and performing SMD+UMD simulations. L747P induces an ordered tight turn in the β3-αC loop, stabilizing the active state over the inactive state by approximately 1.0 kcal/mol (Figure S3C), but not by as large a margin as L747_A750>P. The substantially larger barrier to inactivation in L747_A750>P may result from the proline in its β3αC tight turn coupled with the net three residue deletion (Figure S1B). Altogether, our results suggest that ex19del variants adopt unique conformations near the receiver KD interface that translate into potentially substantial differences in activation propensity.

### L747_A750>P, but not E746_A750 or E746_S752>V, dimerizes in a ligand-independent manner

Previous studies have suggested that KD mutants may promote ligand-dependent “inside-out” dimerization (39). Based on our simulation results, we hypothesized that the L747_A750>P variant forms dimers in the absence of ligand stimulation because it is trapped in a receiver kinase active state. To test our hypothesis, we measured the homo-interaction stoichiometry of each variant in the presence and absence of EGF ligand using two-color pulsed interleaved excitation fluorescence cross-correlation spectroscopy (PIE-FCCS) (33, 40). Live cell PIE-FCCS measurements and analysis were completed on single cells expressing individual ex19del variants with WT data recorded as a negative control for each experiment (see **Methods**).

First, we performed PIE-FCCS experiments in the absence of EGF ligand. Samples were serum starved for 24 hours to ensure no residual ligand-dependent effects. As expected, WT has a median cross-correlation (ƒ_c_) value near zero (ƒ_c_ = 0.01), indicating that it exists predominantly as a monomer. Our results also suggest that E746_A750 and E746_S752>V are predominantly monomeric in the absence of ligand (ƒ_c_ = 0.05 and 0.06, respectively). In contrast, L747_A750>P displays significantly higher median cross-correlation (ƒ_c_ = 0.13) (Figure 3A). Consistent with the cross-correlation values, the diffusion coefficients of eGFP-tagged WT (0.35 μm^2^/s), E746_A750 (0.35 μm^2^/s), and E746_S752>V (0.33 μm^2^/s) are significantly higher than L747_A750>P (0.18 μm^2^/s) (Figure 3B). The increased median cross correlation and decreased diffusion coefficient of L747_A750>P relative to WT is indicative of dimer formation in the absence of ligand stimulation.

**Figure 3.**
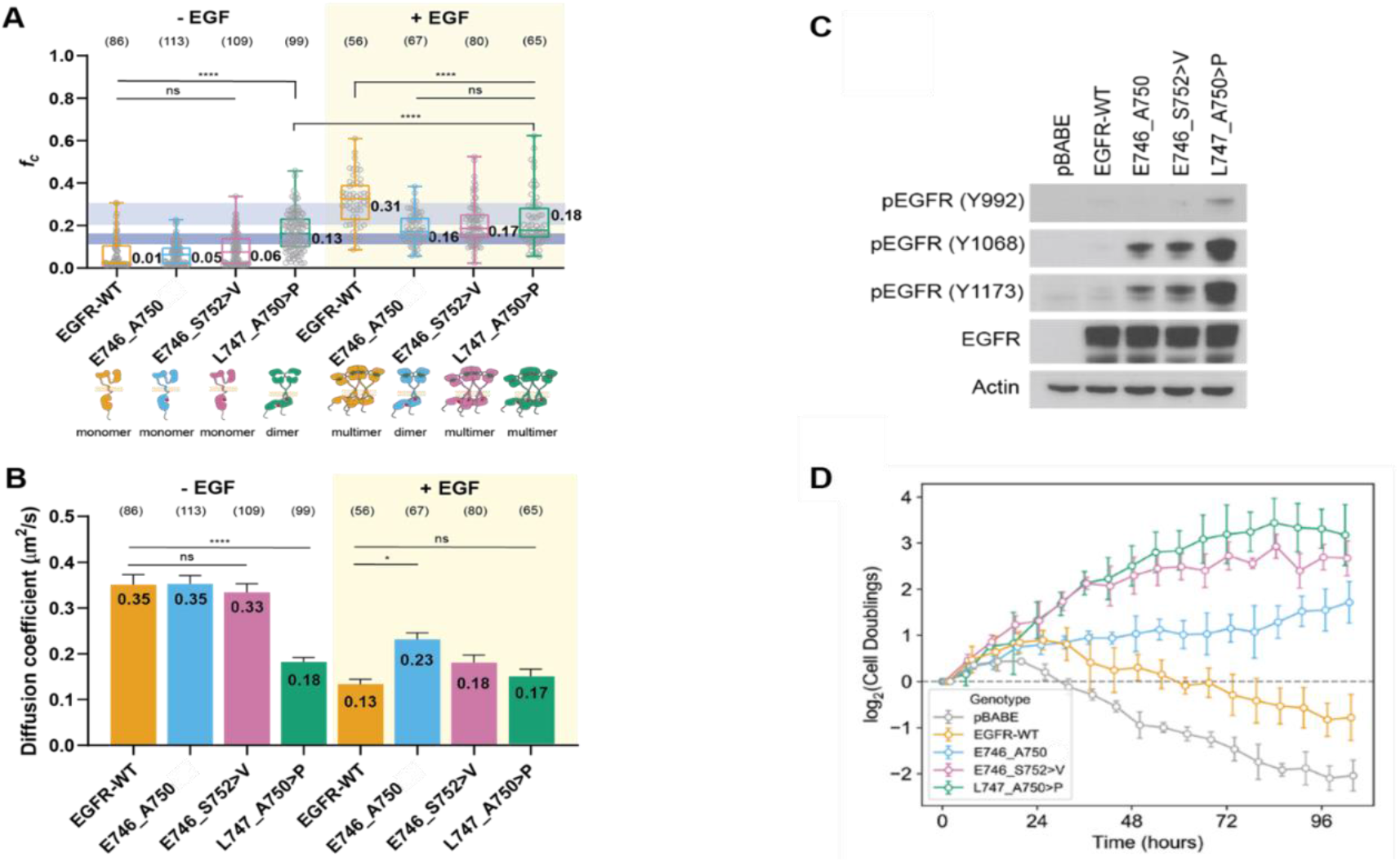
Ex19del variants display allele-specific differences in dimerization and oncogenic growth. **(A)** Cross correlation values of transfected EGFR variants with (+) or without (-) ligand (EGF) stimulation. The dark and light blue boxes indicate the *ƒ_c_* value regions for dimers and multimers, respectively. **(B)** Diffusion coefficient values of EGFR variants with (+) or without (-) ligand (EGF) stimulation. The light orange box indicates EGF-stimulated groups. **(C)** Ba/F3 cells were stably transfected with different EGFR ex19del variants, WT, or empty vector. Cellular lysates were probed with the indicated antibodies to measure phosphorylation. **(D)** Rate of IL-3-independent growth of Ba/F3 cells stably transfected with different ex19del variants, WT, or empty vector.

Next, we performed PIE-FCCS experiments in the presence of EGF ligand to evaluate whether or not ex19del variants differ in their response to extracellular stimulation. A recent study demonstrated that KD mutations can directly change the conformational preferences of the ECD, potentially modulating signaling responses to ligand (41). Here, we observed that WT forms multimers upon stimulation with EGF, consistent with prior studies (ƒ_c_ = 0.31; *D* = 0.13 μm^2^/s) (32, 33, 40, 42). EGF stimulation caused E746_A750 (ƒ_c_ = 0.16; *D* = 0.23 μm^2^/s) E746_S752>V (ƒ_c_ = 0.17; *D* = 0.18 μm^2^/s), and L747_A750>P (ƒ_c_ = 0.18; *D* = 0.17 μm^2^/s) to form a mixture of dimers and multimers (Figure 3A, B). The fact that each of the mutants show lower cross-correlation and faster diffusion compared to WT suggests that the ex19del mutations may have an inhibitory effect on the formation of ligand-dependent multimeric assemblies.

### E746_S752>V and L747_A750>P display enhanced oncogenic activation relative to E746_A750

The strong energetic preference of L747_A750>P to adopt the active conformation (Figure 2E) and corresponding propensity to form ligand-independent dimers (Figure 3A) led us to hypothesize that L747_A750>P would display enhanced oncogenic growth compared with other ex19del variants *in vitro*. To test our hypothesis, we generated expression vectors containing empty vector, WT, E746_A750, E746_S752>V, or L747_A750>P and introduced these into murine lymphoid Ba/F3 cells (43). After selection of stable expression in puromycin, the cells were collected, lysed and blotted for EGFR autophosphorylation (pEGFR). Our results confirmed that all three ex19del variants exhibit strong pEGFR compared to WT. In support of our hypothesis, we observed that L747_A750>P displays substantially higher levels of pEGFR compared with either E746_A750 or E746_S752>V (Figure 3C).

To further investigate ex19del variant differences in IL-3 independent oncogenic growth in Ba/F3 cells, we depleted IL-3 from the growth medium to monitor changes in cell counts over time (Figure 3D). As expected, the Ba/F3 cells expressing either vector or WT EGFR died shortly upon withdrawal of exogenous IL-3, while cells expressing EGFR ex19del variants survived and proliferated. Cells expressing either E746_S752>V or L747_A750>P proliferated at a higher rate compared with cells expressing E746_A750del (Figure 3D). Despite not undergoing ligand-independent dimerization as did L747_A750>P in PIE-FCCS experiments, cells expressing E746_S752>V displayed statistically similar growth rates compared with L747_A750>P. Collectively with our MD simulations, our results suggest that ex19del variants differentially promote growth and enzymatic activity as a function of their energetic barriers to activation.

### E746_S752>V and L747_A750>P are less sensitive to TKI treatment than E746_A750

We considered the possibility that differences may exist between ex19del variant TKI sensitivities, which may explain differences in outcomes between patients with specific ex19dels (4, 19). We previously found that some ex19del variants, in particular E746_S752>V, are especially likely to develop G724S-mediated resistance in response to osimertinib, while L858R and other ex19del variants are not (28, 44). Recently, it was further suggested that L747_A750>P has reduced sensitivity to erlotinib and osimertinib relative to E746_A750 in functional assays due to steric effects (45). Thus, we sought to evaluate the relative TKI sensitivity of E746_A750 in comparison to E746_S752>V and L747_A750>P.

We first treated Ba/F3 cells expressing E746_A750, E746_S752>V, or L747_A50>P with either 30 or 100 nM osimertinib. We observed that autophosphorylation was markedly reduced in both E746_A750 and L747_A750>P, but not in E746_S752>V (Figure 4A). Subsequently, we performed the same experiment in well-established lung adenocarcinoma cell lines expressing E746_A750 (PC9), E746_S752>V (SH450), or L747_A750>P (HCC4006). Again, we observed that E746_S752>V was less sensitive to osimertinib than E746_A750 or L747_A750>P. To model the clinical exposure of EGFR TKIs in lung adenocarcinoma, we performed long-term treatments of osimertinib in these cell lines at a clinically relevant dose (100 nM) (46) with periodic medium/TKI refreshment (Figure 4C). The untreated PC9, SH450, and HCC4006 cells underwent exponential growth and quickly reached confluence within 3 days. The growths of PC9 and HCC4006 cells were inhibited effectively by osimertinib treatment, and the cells initially stopped growing. In particular, the proliferation of PC9 cells was successfully inhibited by osimertinib for more than three weeks. We observed that the HCC4006 cells gradually adapted to the treatment and proliferated to confluence in 20 days. Most notably, however, osimertinib only partially inhibited the proliferation of SH450 cells, and after an incomplete response continued growing, reaching confluence within a week. Thus, consistent with our Western blots, we found that E746_S752>V was least responsive to osimertinib, followed by L747_A750>P, while E746_A750 was completely inhibited (Figure 4C).

**Figure 4.**
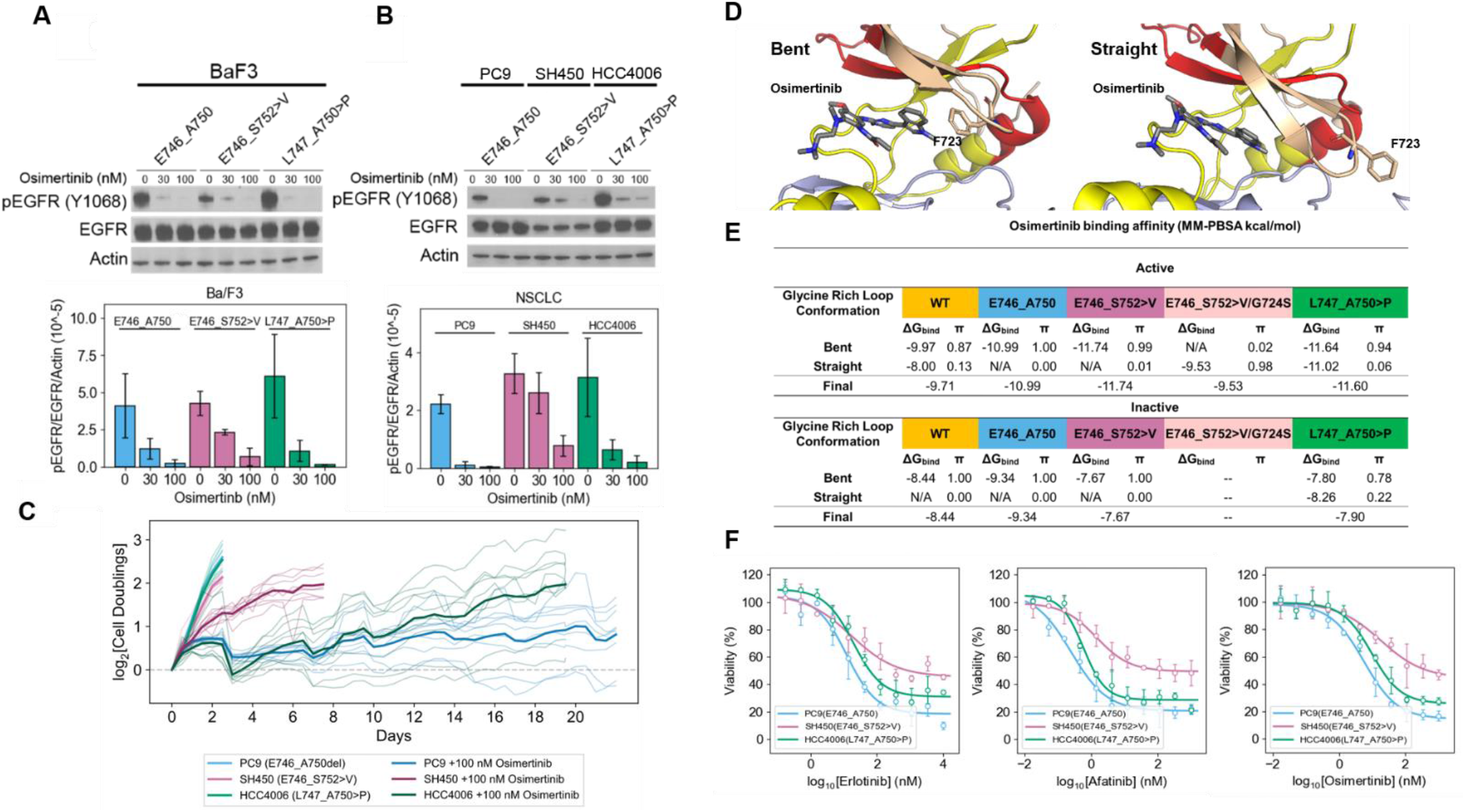
Allele-specific differences in ex19del TKI sensitivity may not be due to differences in TKI binding affinity. **(A)** Ba/F3 cells were stably transfected with different EGFR ex19del variants and treated with increasing concentrations (0, 30, or 100 nM) of osimertinib. Cellular lysates were probed with the indicated antibodies to measure phosphorylation. **(B)** Lung adenocarcinoma cell lines expressing E746_A750 (PC9), E746_S752>V (SH450), or L747_A750>P (HCC4006) were treated with increasing concentrations (0, 30, or 100 nM) of osimertinib. Cellular lysates were probed with the indicated antibodies to measure phosphorylation. Quantifications are represented as the average grayscale ratio of pEGFR/EGFR/Actin+/- standard deviation across three independent biological replicates. **(C)** Time-dependent growth of lung adenocarcinoma cell lines expressing E746_A750 (PC9), E746_S752>V (SH450), or L747_A750>P (HCC4006) treated with either 100 nM osimertinib or buffer. Each condition was performed with 9 replicates (thin lines) and averaged (bold lines). **(D)** Structural models of EGFR in complex with osimertinib in either the bent (F723 facing osimertinib in the ATP binding pocket) or straight (F723 projecting away from the ATP binding pocket) conformations. **(E)** Osimertinib binding affinities for each ex19del variant, WT, and the double mutant E746_S752>V/G724S from simulations starting in the active and inactive states. Bent and straight states were separated by a small 2-state Markov state model based on the G/S724 backbone phi angle. MM-PBSA was not performed if the stationary distribution for a state was estimated at less than 0.05 or the model failed to pass a Chapman-Kalmogorov test. Binding energies are computed as the average MM-PBSA energies of 1000 randomly selected frames from the corresponding MSM cluster. For each EGFR variant, six simulations of 2.0 us each were performed such that there were three each from the active and inactive states (except E746_S752>V/G724S, for which no inactive state simulations were performed). **(F)** Cell viability assays performed in lung adenocarcinoma cell lines stably expressing E746_A750 (PC9), E746_S752>V (SH450), or L747_A750>P (HCC4006) with first (erlotinib), second (afatinib), and third (osimertinib) generation EGFR TKIs.

Based on the *in vitro* data, we hypothesized that E746_S752>V has a lower osimertinib binding affinity than E746_A750 and L747_A750>P. To test this hypothesis, we performed MD simulations of each of the ex19del variants in complex with osimertinib. We performed three independent MD simulations of 2.0 μs each for each EGFR variant (WT, E746_A750, E746_S752>V, E746_S752>V/G724S, or L747_A750>P) bound to osimertinib starting from either the active or inactive conformation (sans inactive E746_S752>V/G724S; 60.0 μs aggregate simulation time). As expected based on the available crystallographic evidence (47), osimertinib binding energies suggested tighter binding in the active state than the inactive state in all cases. Both E746_A750 and L747_A750>P were estimated to have a better osimertinib binding free energy than WT (Figure 4E). Contrary to our hypothesis, E746_S752>V was not predicted to bind osimertinib with a lower affinity than E746_A750 or L747_A750>P. In contrast to previous studies (45), L747_A750>P failed to show a reduced osimertinib binding free energy (Figure 4E).

To better understand our simulation results, we quantitatively evaluated the inhibitory efficacy of three generations of EGFR TKIs (erlotinib, afatinib, and osimertinib) by measuring cell viabilities of isogenic Ba/F3 cells stably transfected with either E746_A750, E746_S752>V, or L747_A750>P in the presence of each TKI separately. We observed that L747_A750>P and E746_S752>V were both at least 10x less sensitive to TKI than E746_A750 (Figure S4). We corroborated these results by measuring cell viabilities of lung adenocarcinoma cell lines expressing different ex19del variants. Here, we also observed that SH450 (E746_S752>V) or HCC4006 (L747_A750>P) were at least 10x less sensitive to erlotinib than PC9 (E746_A750). SH450 were also greater than 10x less sensitive to afatinib and osimertinib as compared to PC9 or HCC4006 (Figure 4F). L747_A750>P displays a similar response to afatinib as E746_A750. Our results suggest that E746_S752>V and L747_A750>P are intrinsically less sensitive to ATP-competitive TKIs *in vitro*. E746_A750 displays the most TKI sensitivity among the three ex19dels.

### Differences in ATP binding may modulate TKI sensitivity across ex19del variants

Our *in vitro* data suggest that E746_S752>V and L747_A750>P display intrinsic resistance to standard first-, second-, and third-generation TKIs. Simultaneously, our MD simulations estimate that E746_S752>V and L747_A750>P reversibly bind osimertinib at least as well as E746_A750. Thus, we hypothesized that the reduced sensitivity of E746_S752>V or L747_A750>P to ATP-competitive inhibitors is the result of higher ATP binding affinities in these receptors than other EGFR oncogenic variants, thereby reducing the relative binding affinity of TKI to ATP.

To test this hypothesis, we estimated the apparent ATP Km and erlotinib Ki for WT, E746_A750, L747_A750>P, and an additional uncommon variant L747_E749 using the ADP-Glo assay as described in the **Methods**. We chose erlotinib for the TKI binding affinity analysis to enable explicit comparison of the effects of ATP Km on noncovalent TKI interactions. Our ADP-Glo assay results suggest that there are substantial differences in ATP kinetics between EGFR variants, consistent previous reports on L858R and G719S (48, 49).

E746_A750 and L747_E749 display ATP Km values of ~100 μM. In contrast, L747_A750>P displays an ATP Km of ~6 μM. Interestingly, the rate of phosphate transfer in L747_A750>P is ~16x lower than E746_A750, but the reduced Km results in comparable catalytic efficiencies (Table 1). In contrast to ATP Km, the difference in erlotinib binding is comparatively small between the tested variants. This results in the apparent erlotinib potency, taken as the ratio of Ki to ATP Km, to be ~6x lower in L747_A750>P than E746_A750 (Table 1). These data are consistent with the reduced sensitivity of L747_A750>P *in vitro* and suggest a general mechanism by which ex19del variants may differ in their responses to TKI.

**Table 1.**
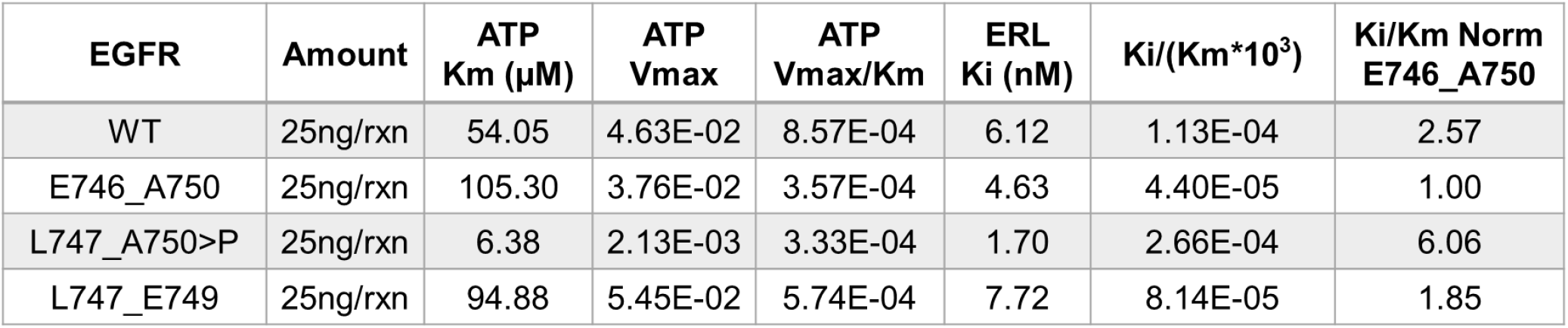
Enzyme kinetic parameters and erlotinib binding affinity for EGFR WT and ex19del variants.

| EGFR | Amount | ATP Km (μM) | ATP Vmax | ATP Vmax/Km | ERL Ki (nM) | Ki/(Km*10<sup>3</sup>) | Ki/Km Norm E746_A750 |
| --- | --- | --- | --- | --- | --- | --- | --- |
| WT | 25ng/rxn | 54.05 | 4.63E-02 | 8.57E-04 | 6.12 | 1.13E-04 | 2.57 |
| E746_A750 | 25ng/rxn | 105.30 | 3.76E-02 | 3.57E-04 | 4.63 | 4.40E-05 | 1.00 |
| L747_A750>P | 25ng/rxn | 6.38 | 2.13E-03 | 3.33E-04 | 1.70 | 2.66E-04 | 6.06 |
| L747_E749 | 25ng/rxn | 94.88 | 5.45E-02 | 5.74E-04 | 7.72 | 8.14E-05 | 1.85 |

Our simulations create structural hypotheses for these differences: First, ex19del variants make distinct hydrogen bonding interactions at the β3αC interface (Figure S5A – D). E746_A750 places S752 at the β3αC *i* + 2 position (Figure 1D) such that the sidechain donates a H-bond to the F723 backbone and is simultaneously stabilized as a H-bond acceptor from the K754 backbone (Figure S5B). Neither E746_S752>V nor L747_A750>P, both of which place a proline at *i* + 2, can make this H-bond (Figure S5C, D). Quantitation of apo-state H-bonding supports this observation, suggesting the glycine-rich loop is more tightly coupled to the β3αC-loop in E746_A750 (Figure S5E). These data, together with previous crystallographic (50) and kinetic (48) studies of EGFR L858R, suggest generally that tight coupling of the β3αC-loop to the glycine-rich loop in αC-helix-stabilizing oncogenic mutants leads to reduced ATP binding affinity.

### New therapeutic strategies may be required to maximally inhibit E746_S752>V-mediated disease

We previously identified the TKI neratinib as a potential therapeutic agent for certain forms of HER2/HER3-mutant cancers in which pan-TKI resistance seems to be associated with enhanced ATP binding affinity (51). Employing the same strategy for neratinib as we did for osimertinib, we performed MD simulations and subsequent MMPBSA binding free energy estimates of ex19dels complexed with neratinib. Our simulations suggest that all of the tested ex19dels reversibly bind neratinib better than osimertinib, but that E746_S752>V has a better neratinib binding energy than E746_A750 or L747_A750>P (Figure 5A). Evaluation of neratinib function inhibition in Ba/F3 cells stably transfected with E746_A750, E746_S752>V, or L747_A750>P demonstrate a complete ablation of pEGFR in E746_S752>V and L747_A750>P at 30 nM. Phosphorylation is largely reduced in E746_A750 at 30 nM and completely ablated at 150 nM (clinical-relevant dose, Figure 5B, C). We also observed that neratinib effectively reduced pEGFR in lung adenocarcinoma cell lines expressing E746_A750, E746_S752>V, or L747_A750>P (Figure 5D – F).

**Figure 5.**
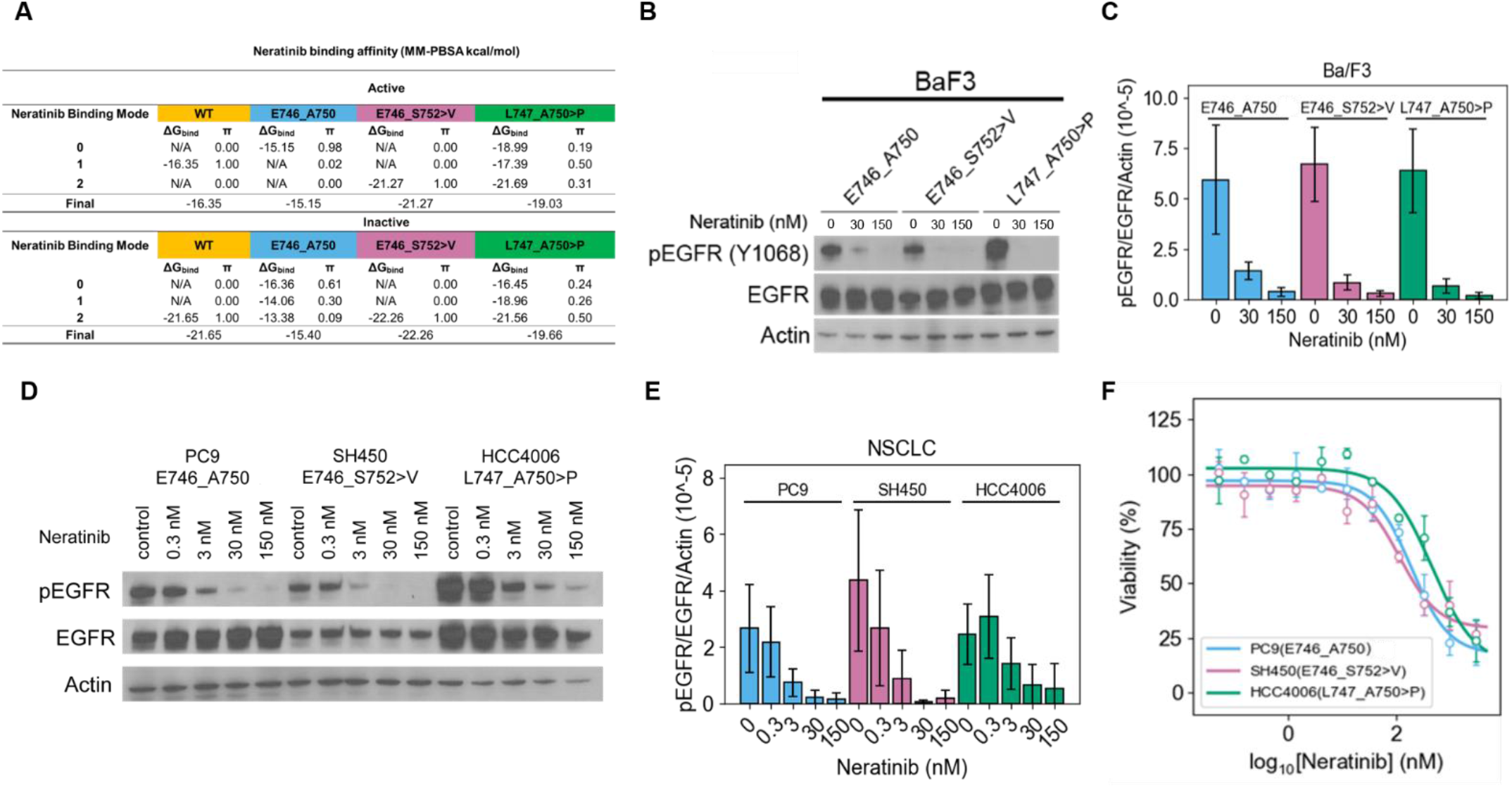
Neratinib effectively inhibits E746_S752>V. **(A)** Neratinib binding affinities for each ex19del variant and WT from simulations starting in the active and inactive states. Three binding modes of neratinib distinguished by the dihedral conformations of the hydroxymethyl pyridine were distinguished with a simple Markov state model. MM-PBSA was not performed if the stationary distribution for a state was estimated at less than 0.05 or the model failed to pass a Chapman-Kalmogorov test for three or two states. Binding energies are computed as the average MM-PBSA energies of 1000 randomly selected frames from the corresponding MSM cluster. For each EGFR variant, six simulations of 2.0 us each were performed such that there were three each from the active and inactive states. **(B)** Ba/F3 cells were stably transfected with different EGFR ex19del variants and treated with increasing concentrations (0, 30, or 150 nM) of neratinib. Cellular lysates were probed with the indicated antibodies to measure phosphorylation. **(C)** Quantification of Ba/F3 neratinib inhibition Western blots are represented as the average grayscale ratio of pEGFR/EGFR/Action+/- standard deviation across three independent biological replicates. **(D)** Ba/F3 cell Lung adenocarcinoma cell lines expressing E746_A750 (PC9), E746_S752>V (SH450), or L747_A750>P (HCC4006) were treated with increasing concentrations (0, 0.3, 3, 30, or 150 nM) of neratinib. Cellular lysates were probed with the indicated antibodies to measure phosphorylation. **(E)** Quantification of lung adenocarcinoma cell line neratinib inhibition Western blots are represented as the average grayscale ratio of pEGFR/EGFR/Actin+/- standard deviation across three independent biological replicates. **(F)** Cell viability assays performed in lung adenocarcinoma cell lines stably expressing E746_A750 (PC9), E746_S752>V (SH450), or L747_A750>P (HCC4006) with neratinib.

## Discussion

Considerable effort has been invested over the last decade to define the molecular mechanisms of oncogenesis and acquired drug resistance in the most commonly occurring *EGFR* mutations, specifically L858R and “exon 19 deletion (ex19del)” (26, 27, 34, 35, 48, 52). These efforts resulted in development of more effective targeted therapies, including today’s first-line therapy for *EGFR*-mutant NSCLC, osimertinib (53). Despite next-generation sequencing having identified the heterogeneity in the various distinct ex19del variants, the allele-specific mechanisms have not been extensively evaluated. The potential reduced likelihood of non-canonical ex19del variants developing T790M or C797S in response to first or third generation TKI, respectively (16, 54), may be because a number of these variants have reduced TKI sensitivity in the setting of higher ATP binding affinity. Indeed, both our group (28) and others (44) found the G724S resistance mutation to occur preferentially to C797S in E746_S752>V and related non-canonical variants in response to osimertinib. However, at present, there has not been a systematic evaluation of patient responses to different TKI based on the specific ex19del variant present in tumor. Thus, it is imperative that we investigate individual ex19del variants pre-clinically to ultimately help guide clinicians in therapeutic decision-making.

Here, we have performed computational, biophysical, and biochemical analyses on a diverse subset of the most frequently occurring ex19del variants: E746_A750, E746_S752>V, and L747_A750>P. Our data show clear differences in the activation profiles and TKI sensitivities of these ex19del variants with potential structural correlates. Specifically, our data suggest that the ligand dependency of receptor activation differs between ex19dels. The L747_A750>P mutant displayed robust αC-helix stabilization from a proline-locked tight turn in MD simulations that translated to ligand-independent dimerization and increased *in vitro* activity in experiments. We also observed that E746_S752>V and L747_A750>P were less sensitive to inhibition by TKI than E746_A750, with E746_S752>V displaying the least sensitivity. We were unable to attribute this effect to binding affinity based on MD simulations of osimertinib or ADP-Glo inhibition assays for erlotinib. Instead, our data suggest a role for variable ATP binding affinity as a potential mediator of these differences in TKI sensitivity. It has previously been observed that some oncogenic EGFR mutations can modulate ATP binding and TKI sensitivity (26, 27, 48, 49).

Collectively, our data demonstrate that ex19dels are a heterogenous group of oncogenic variants. EGFR WT is a monomer in the absence of ligand and stimulated by extracellular EGF to form dimers and multimers/oligomers (Figure 6, yellow). The most frequently occurring ex19del oncogenic mutants, such as E746_A750, increase the propensity for dimerization by stabilizing the acceptor KD (Figure 6, blue). These “classical super acceptors” (35, 52) are ligand-dependent and have lower ATP binding affinity (26), increasing their sensitivity to TKIs with lower reversible binding affinity, such as osimertinib (55). Our simulations and TKI sensitivity data suggest that a subset of ex19del variants, such as E746_S752>V and L747_A750>P, are “tight ATP binders” (Figure 6, pink). These are characterized by ATP binding affinities higher than that of classical super acceptors, making them more resistant to ATP-competitive TKIs, reminiscent of T790M-comutant EGFR (48) (Figure 6). Unlike e.g., L858R/T790M, the apparent inhibitor potency does not differ from the single oncogenic variant e.g., L858R by several orders of magnitude (48); instead, the difference is ~6x. Thus, we distinguish differences in sensitivity from differences in resistance. Finally, another subset of ex19dels, such as L747_A750>P, are characterized by enhanced dimerization propensities greater than that of supper acceptors. These “hyper acceptors” display increased functional activation and exist as ligand-independent dimers (Figure 6, green). The ligand-independent activity of hyper acceptors suggest that some oncogenic variants may be activated via “inside-out” dimerization.

**Figure 6.**
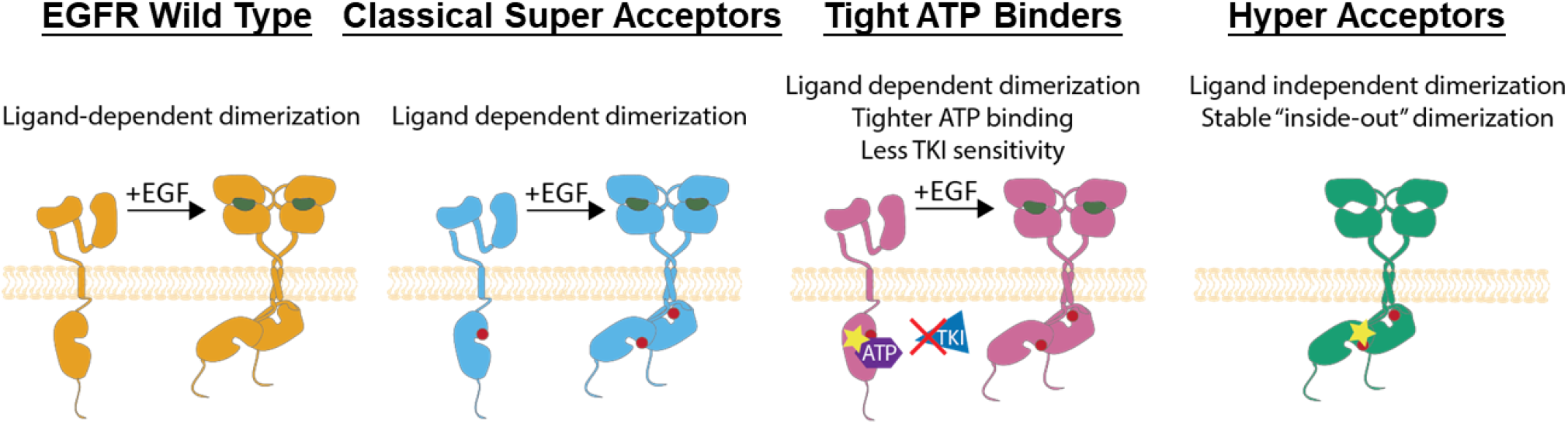
Model of ex19del allele-specific functional differences and strategy for inhibition. **(A)** Discretized classification scheme for EGFR ex19del variants: non-oncogenic with ligand-dependent activation (orange; WT); oncogenic super acceptor with ligand-dependent activation (blue; E746_A750, E746_S752>V); tight ATP binder (pink; E746_S752>V, L747_A750>P); oncogenic hyper acceptor with ligand-independent activation (green; L747_A750>P).

Based on our proposed model, L747_A750>P is both a hyper acceptor and a tight ATP binder, while E746_S752>V is a classical super acceptor and potentially a tight ATP binder. E746_A750 is strictly a classical super acceptor. We hypothesize that ex19del variants exist along a spectrum of dimerization propensities and ATP affinities. Based on predicted structural similarity to the mutants studied in depth here, we propose initial classifications of the rarer ex19del variants identified in AACR GENIE along this spectrum. We anticipate that additional functional characterization of ex19del variants along these axes will allow more personalized treatment of ex19del NSCLC patients.

Generally, our data lead us to suggest that treatment of ex19del variants may require unique consideration of the variant’s functional properties. For example, we speculate that mutations with enhanced ligand-independent dimerization would be less amenable to EGF-blocking antibody / TKI combination therapies than classical super acceptor-like variants. We also suggest that for ex19dels with high ATP binding affinities, the use of covalent TKIs with higher reversible binding affinities may be necessary to overcome reduced TKI sensitivity, such as neratinib or mobocertinib. Alternatively, because increasing the reversible binding affinity on covalent inhibitors can reduce mutant selectivity and cause undesirable side-effects, recognition of tight ATP binding ex19dels may motivate the design of PROTAC-based or allosteric inhibitors. To facilitate future structural comparisons of Ex19Del variants, we have made our computational structural models of these variants available on GitHub.

This study is not a comprehensive guide to EGFR ex19del variants. We hope that subsequent work expands upon this study to better characterize uncommon ex19dels. While *in silico* modeling can provide useful insight to generate hypotheses, it can be limited by factors such as the quality of the predicted structures, the short simulation timescales currently accessible, the start- and end-state dependency of umbrella sampling simulations, and the simplification of the system from transmembrane dimers/multimers to monomeric intracellular KDs. Similarly, *in vitro* data in the absence of structural characterization and dynamical insight can make it challenging to generalize findings and perform rational drug design. We anticipate that continued characterization of ex19del structures through experimental structural biology, detailed kinetics studies, and receptor signaling/crosstalk studies will be an important next step in ongoing efforts to design new treatment strategies for patients with EGFR-mutant NSCLC.

## Materials and Methods

### Tyrosine kinase inhibitor source and preparation

Inhibitors were purchased from Selleck Chemicals.

### Cell culture

Ba/F3 cells (DSMZ), PC9 (ATCC), SH450 (ATCC), and HCC4006 (ATCC) were cultured in RPMI 1640 with L-glutamine (Mediatech) supplemented with 10% heat-inactivated FBS (Thermo Fisher Scientific), penicillin (100 U/mL; Thermo Fisher Scientific), streptomycin (100 μg/mL; Thermo Fisher Scientific), and IL3 (1 ng/mL; Thermo Fisher Scientific) until retroviral transduction and subsequent IL3 withdrawal. Cells were grown in a humidified incubator with 5% CO_2_ supply at 37°C. Mycoplasma contamination was evaluated routinely during cell culture using a VenorGeM Mycoplasma Detection Kit (Sigma-Aldrich).

### Generation of EGFR-expression constructs and generation of Ba/F3 cell lines

pBabe plasmids with EGFR ex19del mutation encoding cDNAs (EGFR E746_A750, EGFR E746_S752>V, EGFR L747_A750>P) and EGFR WT were purchased from Addgene. The empty pBABE-puro retroviral vector or pBABE-EGFR mutants were transfected, along with the envelope plasmid pCMV-VSV-G (Cell Biolabs, San Diego, CA, USA), into cells Plat-GP packaging cells (Cell Biolabs). 48 hours after transfection, viral media was collected, and the debris were removed by centrifugation. For each separate transduction, 1 × 10^6^ Ba/F3 were re-suspended in the viral media and supplemented with 10 μg/mL polybrene (Santa Cruz Biotechnology, Dallas, TX, USA). Transduced cells were selected using 2 μg/mL puromycin (Invitrogen). EGFR construct expressions were checked before experiments, and only stable polyclonal populations were used.

### Quantitative assessment of cell proliferation during IL-3 withdrawal

Ba/F3 cells that had been transduced with EGFR-expressing constructs, selected with 2 μg/mL puromycin, and growing in media containing 1 ng/mL IL-3 were washed twice with warm PBS to remove IL-3. Cells were re-suspended in media without IL-3 and seeded in 96-well imaging plates at a density of 3,000 cells/well. Cells were periodically scanned in IncuCyte® ZOOM every 6 hours using Incucyte® Nuclight Rapid Red Dye for nuclear labeling. Cell doubling values were calculated using the cell counts at each time point divided by the cell counts at start time point.

### Immunoblot and antibodies

Antibody EGFR (#2232), pEGFR Y1068, pEGFR Y992, pEGFR Y1184, horseradish peroxidase (HRP)-conjugated anti-rabbit (#7074) were all purchased from Cell Signaling Technology, and the actin antibody (A2066) was purchased from Sigma-Aldrich. For immunoblotting, cells were harvested before or after ligand or drug treatment, washed using PBS, and lysed with RIPA buffer [50 mmol/L Tris HCl (pH 8.0), 150 mmol/L sodium chloride, 5 mmol/L magnesium chloride, 1% Triton X-100, 0.5% sodium deoxycholate, 0.1% SDS, 40 mmol/L sodium fluoride, 1 mmol/L sodium orthovanadate, and complete protease inhibitiors (Roche Diagnostics)]. For signal detection, Western Lightning ECL reagent (PerkinElmer) was used. Phosphorylated bands were quantified using ImageJ.

### Viability assays

Experiments were conducted in the Vanderbilt High Throughput Screening Facility. Cells were seeded at approximately 800 cells per well in 384-well plates using Multidrop™ Combi Reagent Dispenser (Thermo Scientific). Medium containing different drug concentrations were prepared using a column-wise serial 3X dilution in 384-well plates using a Bravo Liquid Handling System (Agilent) and were added to the cells. Cell viabilities are obtained using CellTiter-Blue® Cell Viability Assay (Promega).

### Statistical analysis

All experiments were performed at least three time and the difference were determined by ordinary one-way ANOVA using GraphPad Prism 9.2.0. Difference was considered significant when *p* < 0.05.

### Enzymatic analysis

EGFR WT (#E10-112G, lot J3837-8), E746_A750 (#E10-122JG, lot O3886-10), L747_A750>P (#E10-12MG, lot G1200-3), and L747_E749 (#E10-12LG, lot G1344-5) were purchased from SignalChem. The Promega ADP-Glo™ kinase assay kit was used to quantify the amount of ADP produced by each EGFR variant in 1XBFA buffer and in the presence or absence of erlotinib at varying concentrations. Poly(4:1 Glu, Tyr) at a concentration of 0.2 μM was used as the peptide substrate. Reactions were performed at room temperature for 40 minutes each at varying ATP concentrations: 3.125, 6.25, 12.5, 25, 100, 500 μM. Reactions were performed on 384-well plates with each ATP concentration performed in duplicate. Following incubation for 40 minutes, the Promega ADP-Glo™ reagent is utilized to quench the enzymatic reaction and remove residual ATP. The kinase detection agent provided with the assay kit is subsequently used to convert product ADP back into ATP and measure luminescence from the ATP-powered luciferase/luciferin reaction. ATP Km and erlotinib Ki were fit according a mixed model of inhibition using GraphPad Prism 9.3.1.

### Pulsed Interleaved Excitation Fluorescence Cross-Correlation Spectroscopy (PIE-FCCS)

FCCS data were taken on a customized microscope system to introduce pulsed interleaved excitation (PIE) and time-correlated single photon detection as shown in previous works. (33, 56). A supercontinuum pulsed white laser (9.74 MHz repetition rate, SuperK EXW-12 NKT Photonics, Birkerød, Denmark) was split into 488 nm and 561 nm using filters and mirrors for the excitation of eGFP and mCherry, respectively. The 50 ns time delay for PIE was introduced by directing the splitted beams through two different-length optical fibers (57). The beams were cleaned, overlapped, and directed to the microscope. A 100X TIRF oil objective (Nikon, Tokyo, Japan) was used for the excitation beam focus and fluorescence emission collection. NIST traceable fluorescein (50 nM; Thermo Fisher Scientific) was used for optical path alignment, and a short, fluorescent-tagged DNA was used as both alignment and as *ƒ_c_* value control. Previously published negative and/or positive controls (33, 56) were tested before the experiment for data quality control and comparisons of the fit parameters. The overlapped excitation beams were focused on to the fluorescently tagged EGFR (WT or mutant)-transfected COS7 cell membrane. The z axis scan was done to ensure that the laser beam was focused on the flat, peripheral membrane area. One 60-second data acquisition was taken per area per cell. The emitted fluorescence was collimated, separated, and filtered before focused onto single-photon avalanche diodes (Micro Photon Devices, Bolzano, Italy) independently. A time-correlated single photon counting module (Picoharp 300, PicoQuant, Berlin, Germany) recorded the time-tagged photon counts for each channel. For analysis, the time-tagged photon counts were divided into six 10-second acquisitions, binned, and gated for channel differentiation. Auto- and cross-correlation curves corresponding to each species were calculated and generated using a custom MATLAB script. Curves of each acquisition per area were filtered, averaged, then fitted to a single component, 2D diffusion model as shown previously (56–58). The averaged and fitted auto-correlation curves show the average dwell time (τ_D_) that we use to calculate the effective diffusion coefficient, D_eff_ = ω_o_^2^/4τ_D_. The amplitude of the curves can be used to calculate the local concentration of the diffusing receptors in the detection area. Using the cross-correlation curve, we can calculate cross-correlation values (*ƒ_c_*) that indicate the degree of oligomerization. Based on the *ƒ_c_* calibration using live cell control system, expected *ƒ_c_* value for a monomer-dimer equilibrium is 0.10 to 0.15. Higher *ƒ_c_* values indicates higher order oligomerization (57).

### Computational modeling

Structural modeling of proteins was carried out using the Rosetta v.3.12 package (59, 60). Molecular dynamics simulations were performed with Amber18 utilizing the Amber ff14SB and GAFF2 forcefields for proteins and ligands, respectively (51, 61). We estimated protein-ligand binding free energies using the MMPBSA.py package in AmberTools18 (62). RMSD, atom-atom distances, and dihedrals angles were obtained using CPPTRAJ in AmberTools18. Markov modeling analysis was performed with PyEMMA2 (63). The initial structure of osimertinib was taken from PDB ID 4ZAU (47). The initial structure of neratinib was obtained PDB ID 3W2Q (64). The structures were geometry optimized using Gaussian 09 revision D.01 at B3LYP/6-31G(d) level of theory and the electrostatic potential of the optimized structures computed with HF/6-31G(d) in the gas phase. Atomic partial charges were fit with the restrained electrostatic potential (RESP) algorithm in AmberTools18. ATP parameters were developed previously (65) and coordinates initialized from PDB ID 2ITX. For protein-ligand complexes of variants with osimertinib, neratinib, or ATP, we utilized the above PDB structures for ligand placement.

### EGFR ex19del structural modeling

We first built structural models of the 60 ex19del variants identified in AACR GENIE with RosettaCM using the REF2015 score function(66). As templates, we selected the active state EGFR WT structures from PDB IDs 2ITX and 2GS6. We also used the active state model of L858R from PDB ID 4I20. We also included as templates the MD equilibrated structural models of E746_A750 and E746_S752>V we made for our prior study (28). We generated 5,000 RosettaCM models for each variant. The best scoring variant from each was simulated with GaMD for 1.0 us (60.0 us total). GaMD simulation trajectories were clustered with DBSCAN in CPPTRAJ based on β3αC loop RMSD. Each variant was subsequently remodeled with RosettaCM to generate 10,000 more models using the DBSCAN cluster centers as additional templates alongside the prior templates. The best scoring model in round two is the final model. Active state L747P was modeled as a point mutation using the Rosetta PackRotamersMover and FastRelax mover starting from EGFR WT in PDB ID 2ITX. We performed a 1.0 us GaMD simulation on the resulting L747P structure, followed by DBSCAN clustering with CPPTRAJ as above. A representative structure from each cluster was relaxed in Rosetta with progressively ramped-down constraints to the starting coordinates to produce 50 models for each cluster. The best scoring model was carried forward for additional simulations. Inactive state structural models of E746_A750, E746_S752>V, L747_A750>P, L747_T751, L747_P753>S, and L747P were modeled with RosettaCM using the inactive state symmetric dimer EGFR WT in PDB ID 3GT8 as a template.

### Conventional MD (cMD) simulations

Each structure was solvated in a rectangular TIP3P box (12 Å buffer) neutralized with monovalent Cl^-^ and Na^+^ ions (49). Minimization proceeded in three stages: solvent minimization with constraints on solute atoms, solute minimization with constraints on solvent, and subsequently full system minimization without constraints. Each of these stages consisted of 1,000 steps of steepest gradient descent followed by 4,000 steps of conjugate gradient descent. The system was heated in the canonical (NVT) ensemble to 100 K over 100 ps. The system was then heated in the isothermal-isobaric (NPT) ensemble at 1 bar from 100 K to physiologic 310 K over 400 ps. Equilibration was performed in NPT ensemble at 310K for an additional 1000 ps. NPT simulations utilized a Monte Carlo barostat. The temperature was controlled using Langevin dynamics with a collision frequency of 2.0 ps^−1^. A unique random seed was used for each simulation. SHAKE was implemented to constrain bonds involving hydrogen atoms. Periodic boundary conditions were applied and the particle mesh Ewald (PME) algorithm was adopted for long-range electrostatics with a switching distance of 10 Å. Hydrogen mass repartitioning was employed on solute atoms to allow an integration time step of 4 fs.

### Gaussian Accelerated MD (GaMD) simulations

Gaussian accelerated MD (GaMD) is an enhanced sampling method that adds a boost potential to the potential energy surface to accelerate transitions between low-energy states (51, 67). The dual boost potential scheme was applied to the system in order to enhance conformational sampling (67). Systems were equilibrated for 50 ns in cMD. Subsequently, potential statistics for GaMD acceleration were computed from a 10 ns cMD simulation. After addition of the GaMD boost potential, simulations were equilibrated for an additional 50 ns before production. All GaMD simulations were performed in NVT ensemble with a Langevin thermostat and collision frequency of 5.0 ps^−1^. The upper limit of the boost potential standard deviation was set to 6.0 kcal/mol.

### Umbrella sampling and conformational free energy landscapes

Conformational free energy landscapes (FEL) of EGFR WT and ex91del mutants were obtained with constant velocity steered MD (SMD) coupled with Umbrella sampling (US) simulations. The weighted histogram analysis method (WHAM) as implemented by Alan Grossfield (15) was used to perform final statistical reweighting of the US simulations. SMD simulations of 100 ns were performed with a harmonic bias potential and spring constant of 1000 kcal/mol/Å^2^. SMD simulations were performed from the active to the inactive state and vice versa using Cα RMSD to the reference coordinates as the collective variable. A minimum of 250 windows were selected from each forward and backward simulation with which to seed US simulations. Therefore, a total of at least 500 windows per system were used to ensure overlap. A 2D harmonic restraining potential was applied to two CVs for the US simulations. CV1 (y-axis) was defined as the difference in the distance between K860(NZ) – E762(OE1, OE2) and K745(NZ) – E762(OE1, OE2). CV2 (x-axis) was defined as the dihedral angle formed by the Cα atoms of the following residues: D855, F856, G857, and L858. A 2.0 kcal/mol/Å^2^ spring constant was used for CV1, and a 10.0 kcal/mol/rad^2^ spring constant was used for CV2. At each umbrella center a 5 ns simulation was performed. The first 1 ns was used for equilibration, and the following 4 ns were used for analysis in WHAM. Lowest free energy pathway (LFEP) analysis completed with the LFEP package freely available from the Moradi Laboratory at the University of Arkansas.

### Markov model analysis of molecular dynamics simulations

We constructed hidden Markov state models (MSM) to distinguish between two backbone conformations of the glycine-rich loop at residue positions 723 and 724 for osimertinib binding free energy estimates. We also constructed MSMs to distinguish between up to three dihedral conformations of the hydroxymethyl pyridine ring of neratinib for binding free energy estimates. Each MSM was constructed with 6.0 μs (3 × 2.0 μs for each variant for a given active/inactive state) of MD simulation trajectories where frames were collected every 100 ps. All MSMs were constructed with a lag time of 100 ps. The discretized feature trajectories were clustered using KMeans clustering into 500 microstates. All MSMs were validated with Chapman-Kolmogorov tests. In the case of the neratinib binding mode MSMs, if a receptor-neratinib complex did not sample three binding modes, the MSM was regenerated as a two-state model. If only a single dihedral conformer was effectively sampled throughout all three simulations, it was manually assigned a stationary distribution of 1.0. Otherwise, stationary distributions were estimated from MSMs and used to weight the estimated non-covalent binding free energy.

### Binding free energy calculations

The estimated binding free energies between EGFR and TKI (osimertinib or neratinib) was computed with the molecular mechanics Poisson-Boltzmann surface area (MM-PBSA) method using the MMPBSA.py program in AmberTools18 (62). From each MSM metastable state, we randomly resampled 1,000 structures to use for binding free energy calculations. For the MM-PBSA calculations, the internal and external dielectric constants were set to 4.0 and 80.0, respectively. The nonpolar component of the solvation free energy was estimated from the solvent accessible surface area with the classical method (INP=1) using default coefficient and offset values. Atomic radii were taken from the parameter-topology file (RADIOPT=0).

## Acknowledgments

This work was supported by NIH NCI R01CA227833 and NIH NIGMS R01 GM080403. CML was also supported in part by EGFR RESISTERS/LUNGEVITY Lung Cancer Research Award, P30-CA086485, UG1CA233259, and 5P01CA129243-12. ZD is supported by the 2018 AACR-AstraZeneca Lung Cancer Research Fellowship, Grant Number 18-40-12-DU. Work in the Meiler laboratory is supported through the NIH (R01GM080403, R01GM099842, and R01GM073151). AWS and SK were supported by the National Science Foundation (CHE-1753060). BPB is supported through the NIH by a Ruth L. Kirschstein NRSA fellowship (F30DK118774).

## Data availability

Data are available upon request. Computational structural models for EGFR ex19del active state kinase domains are available on the Meiler Lab GitHub repository. Example scripts for each computational modeling section are also available on the Meiler Lab GitHub repository. Please contact the corresponding authors with additional questions.

## Abbreviations

AACR: (American Association for Cancer Research)
GENIE: (Genomics Evidence Neoplasia Information Exchange)
ex19del: (exon 19 deletion)
MD: (molecular dynamics)
Rosetta: 
PIE-FCCS: (pulsed interleaved excitation fluorescence cross-correlation spectroscopy)
TKI: (tyrosine kinase inhibitors)

---

**Figure S1.**
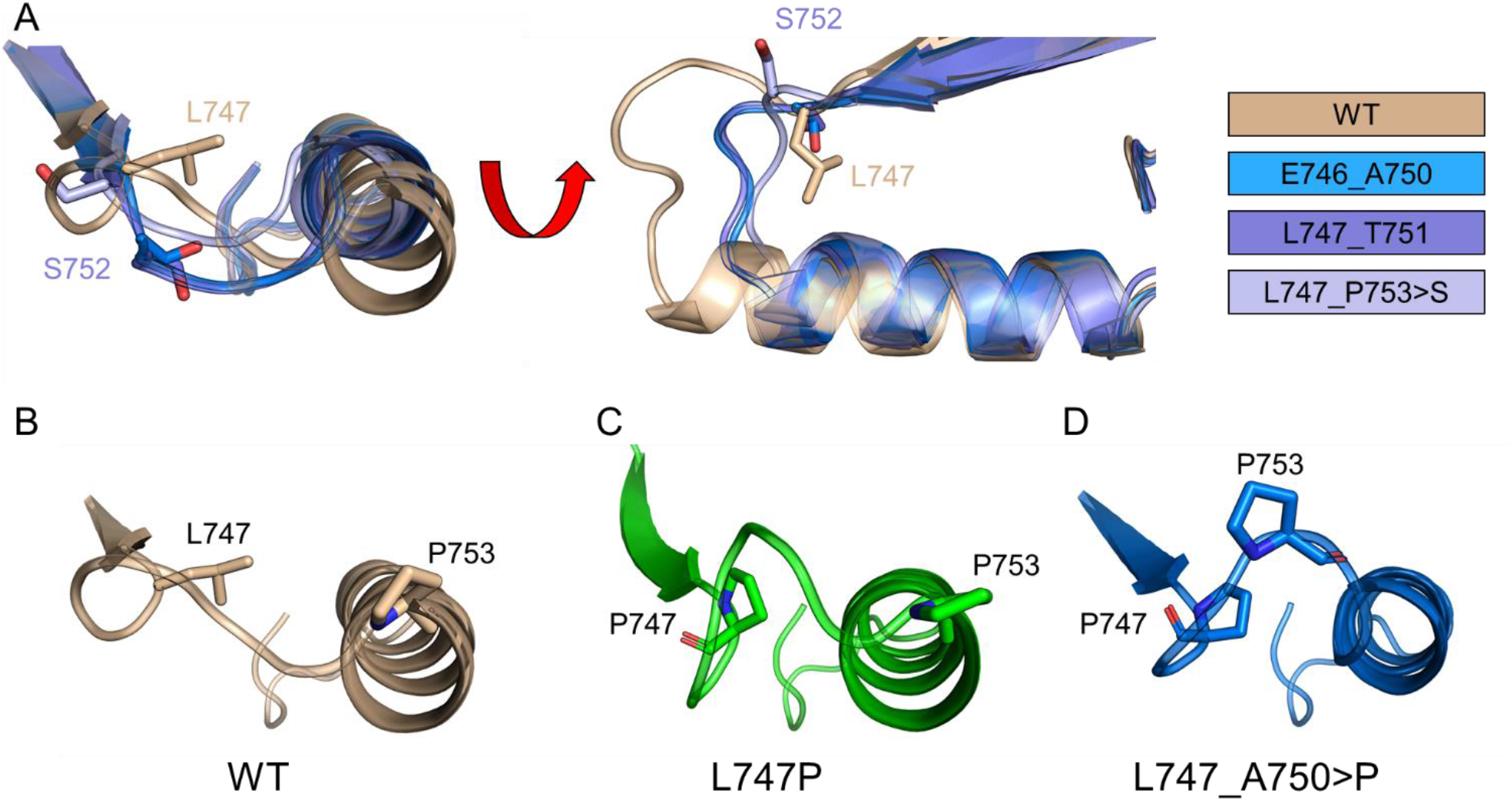
Structural comparison of modeled ex19del β3αC motifs. **(A)** Superimposition of the β3αC region of the most common ex19del variants with WT. Rendering of the β3αC loop in **(B)** WT, **(C)** L747P, and **(D)** L747_A750>P. L747P and L747_A750>P both form a tight turn in the β3αC loop. The L747_A750>P tight turn contains a proline in the second position and fewer residues on the N-terminus of the αC-helix.

**Figure S2.**
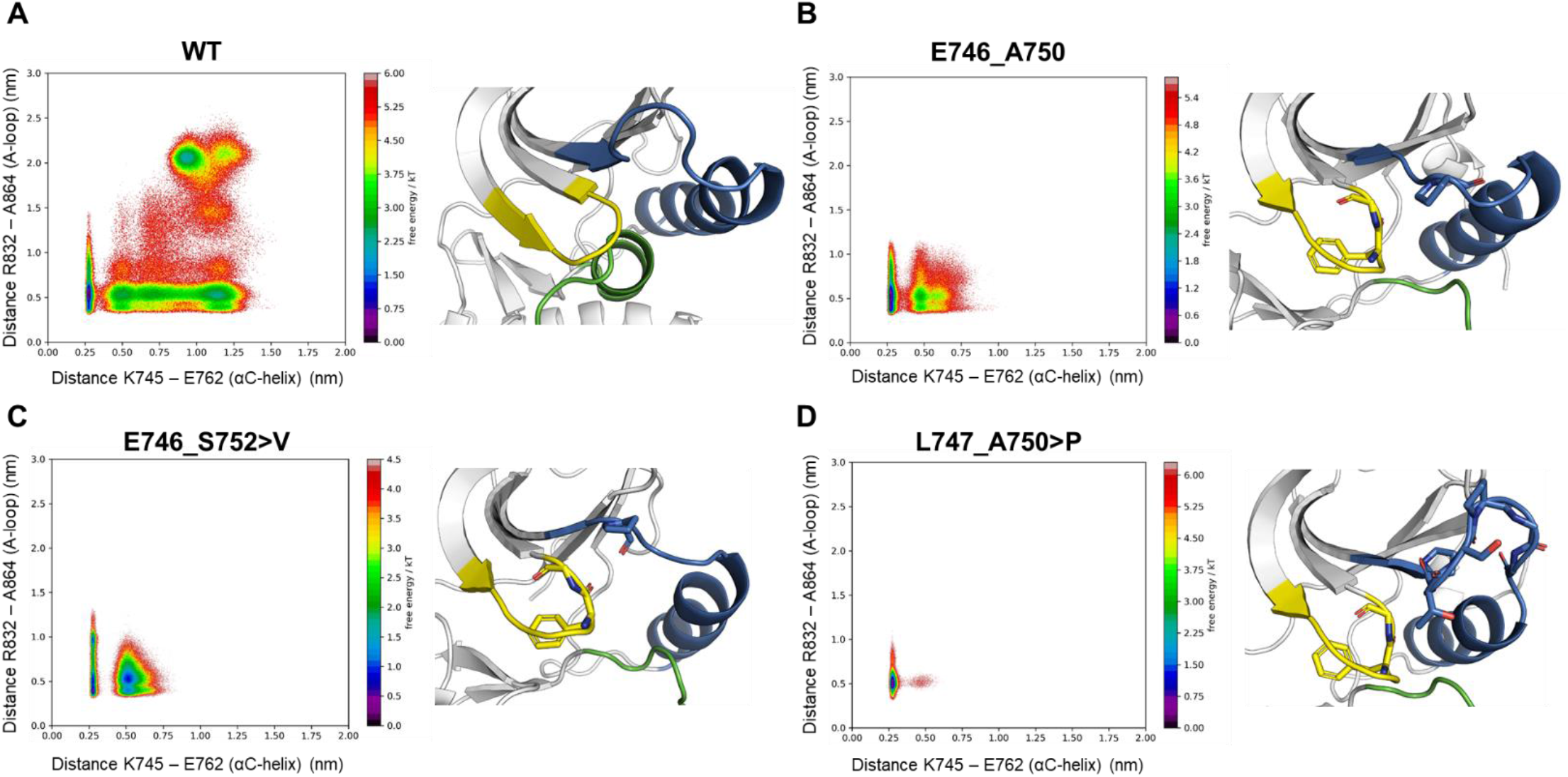
Conventional MD simulations of several ex19del variants starting from the active state. Boltzmann-weighted probability distributions of **(A)** WT, **(B)** E746_A750, **(C)** E746_S752>V, and **(D)** L747_A750>P conformational changes in conventional MD simulations. All simulations were started from the active state. Three independent simulations for each system were run for 4.0 us each. The inward/outward motion of the activation loop is depicted on the y-axis (larger numbers indicate more inward), and the inward/outward motion of the αC-helix is depicted on the x-axis (larger numbers indicate more outward). Snapshots are from the end of one of the three independent simulations. WT transitioned to the Src-like inactive state in one of the three simulations. The glycine-rich loop is colored yellow, the β3αC-loop and αC-helix are blue, and the activation loop is green.

**Figure S3.**
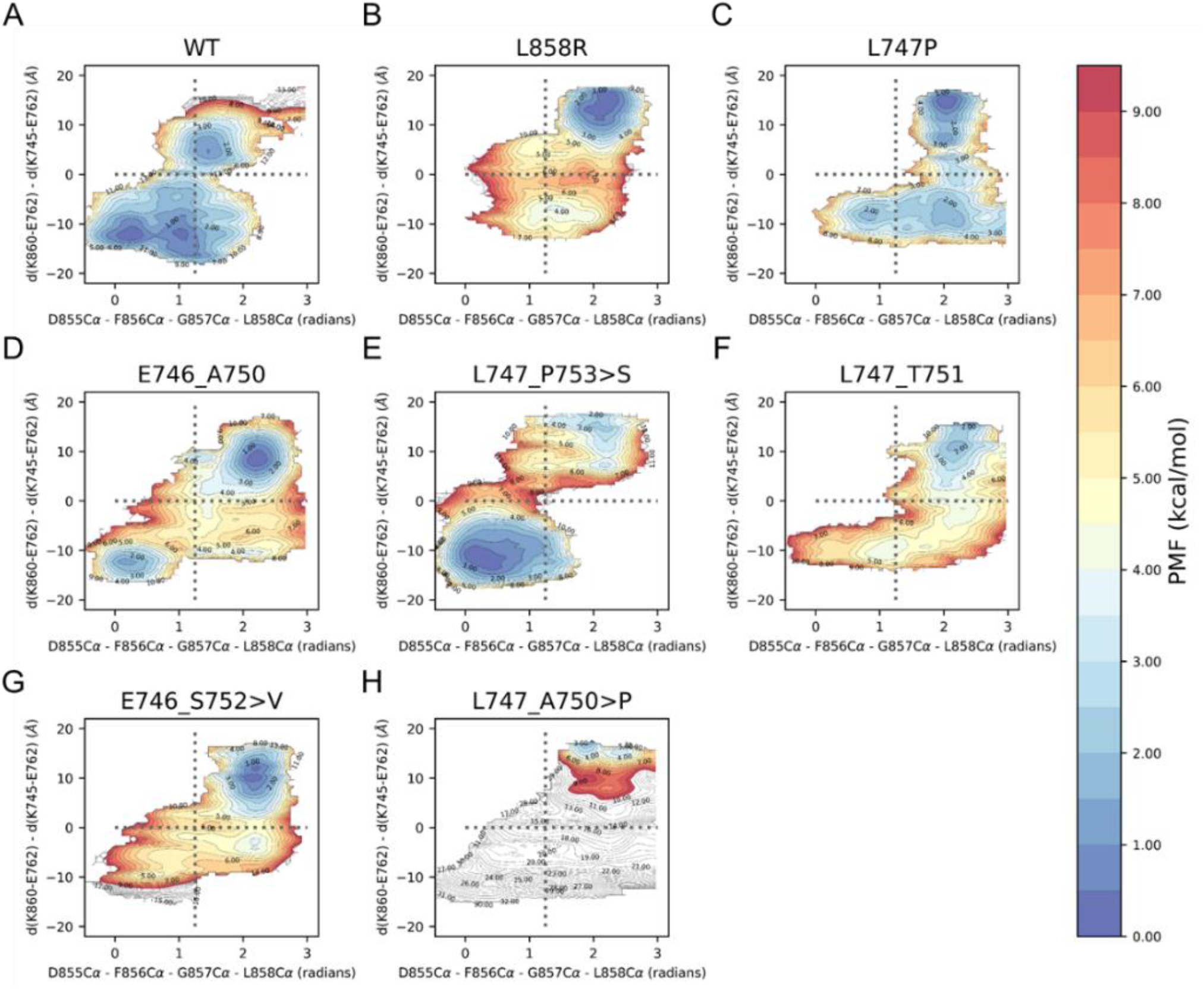
Conformational free energy landscapes of EGFR variants from umbrella sampling MD simulations. Collective variables describe the active and inactive states as the pseudo-dihedral angle formed by the alpha carbon atoms of residues D855, F856, G857, and L858 (x-axis) as well as the difference in distance between the capping sidechain atoms of E762 and K745 (d1) and E762 and K860 (d2) (y-axis). Conformational free energies are shown for **(A)** WT, **(B)** L858R, **(C)** L747P, **(D)** E746_A750, **(E)** L747_P753>S, **(F)** L747_T751, **(G)** E746_S753>V, and **(H)** L747_A750>P. Plots are contoured at 0.5 kcal/mol and colored within the range 0 (blue) and 9.5 (red) kcal/mol. Contours above 9.5 kcal/mol are colored white.

**Figure S4.**
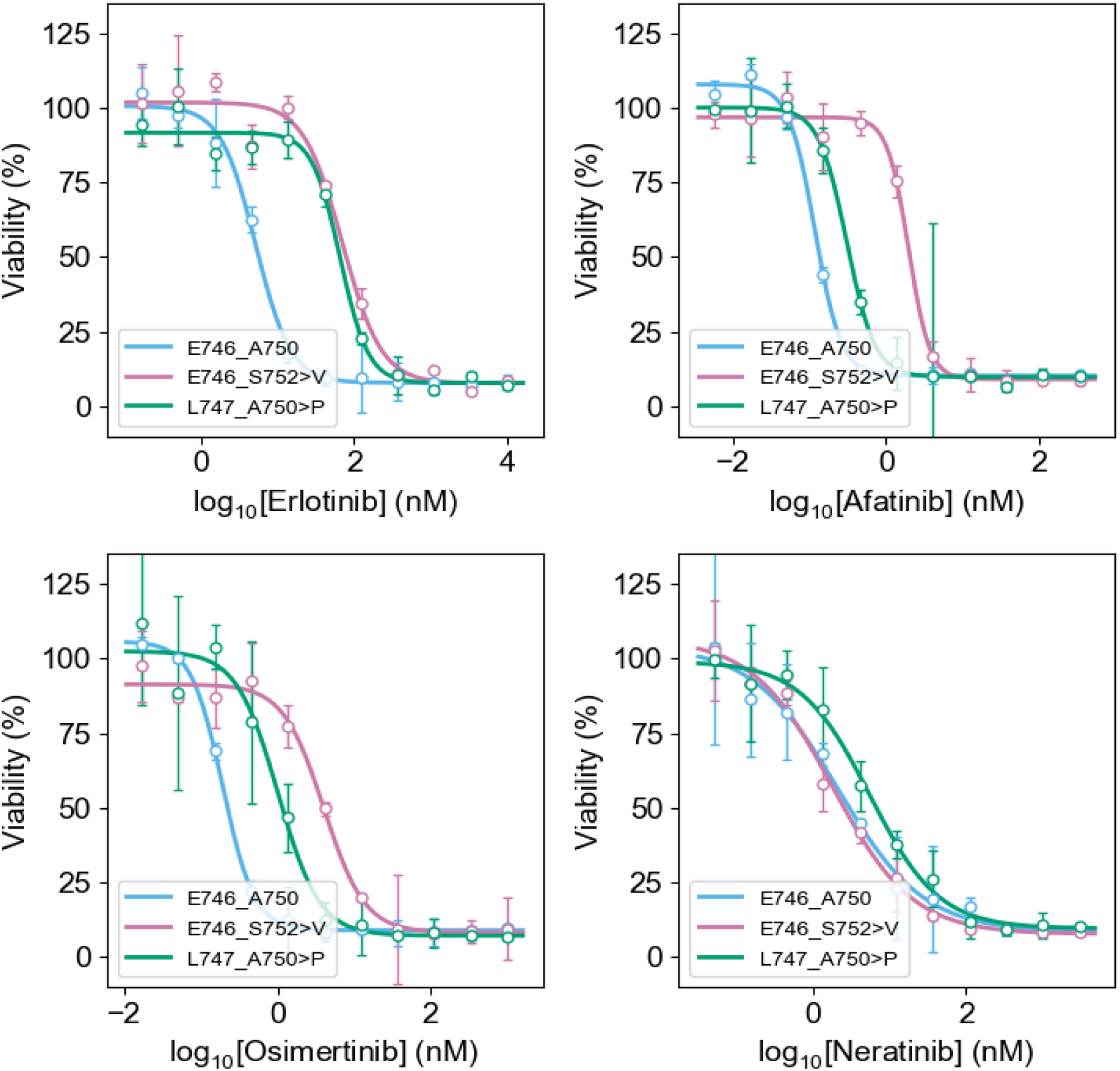
TKI sensitivities of Ba/F3 cells expressing EGFR mutants. Cell viability assays performed in Ba/F3 cells stably expressing E746_A750 (blue), E746_S752>V (pink), or L747_A750>P (green) with erlotinib, afatinib, osimertinib, or neratinib.

**Figure S5.**
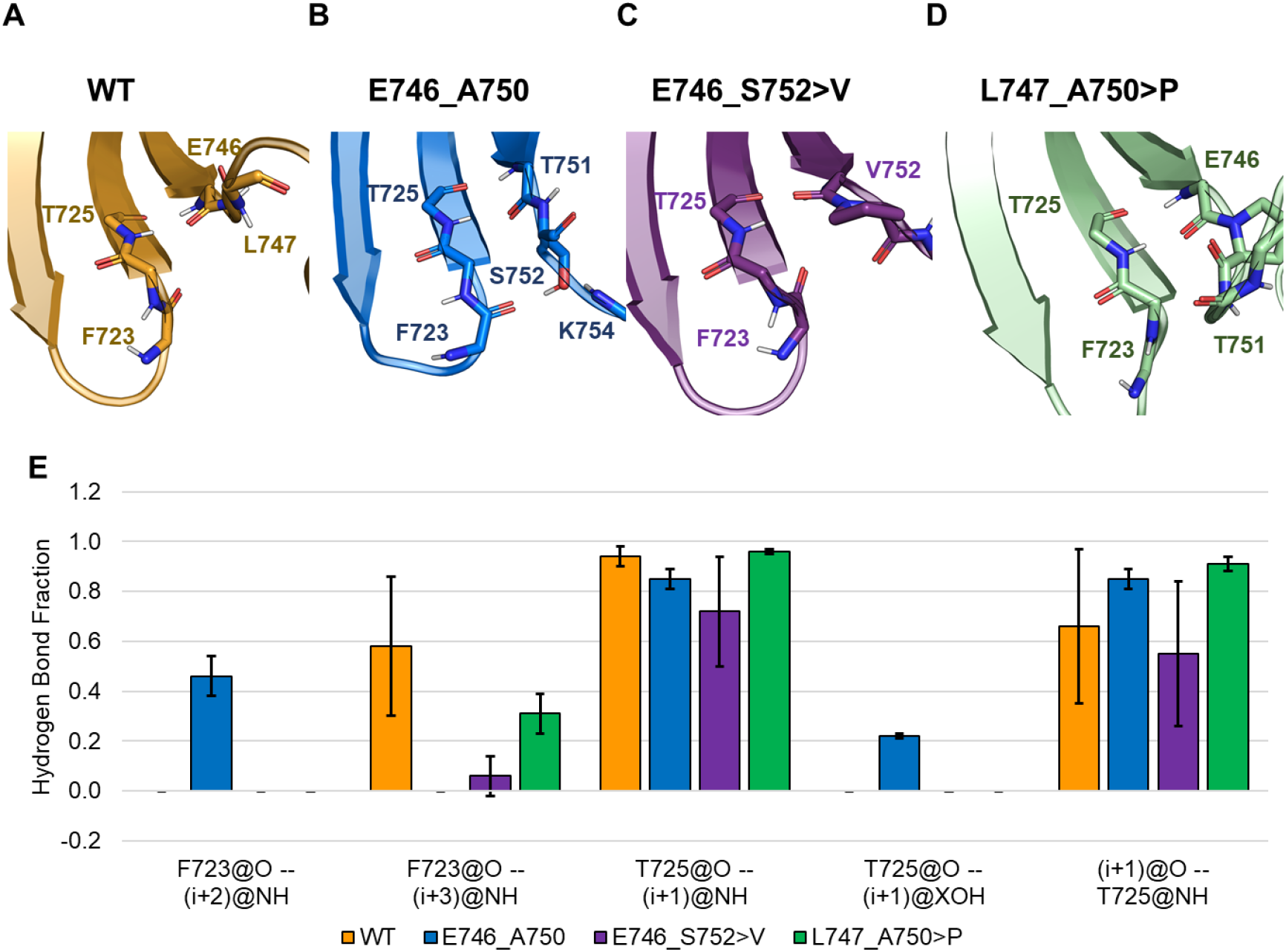
Conventional MD simulations demonstrate ex19del β3αC hydrogen bond networks. Apo-state conventional MD simulation snapshots of β3αC hydrogen bond networks in **(A)** WT, **(B)** E746_A750, **(C)** E746_S752>V, and **(D)** L747_A750>P. **(E)** Quantification of hydrogen bond stability of select β3αC hydrogen bonds at the interface. Hydrogen bonds are defined by donor/acceptor heavy atom distances of ≤ 3.5 and angles between 135 and 180 degrees. Quantifications are based on three independent trials of 4.0 us apo-state simulations of each system starting from the active state.

**Table S1.** Exon 19 deletion variants identified in AACR GENIE database.

| Mutation | # Cases | Frequency (%) |
| --- | --- | --- |
| p.E746_A750del | 857 | 65.07 |
| p.L747_P753delinsS | 101 | 7.67 |
| p.L747_T751del | 71 | 5.39 |
| p.E746_S752delinsV | 54 | 4.10 |
| p.L747_A750delinsP | 51 | 3.87 |
| p.L747_E749del | 24 | 1.82 |
| p.S752_I759del | 18 | 1.37 |
| p.E746_T751delinsA | 17 | 1.29 |
| p.L747_S752del | 14 | 1.06 |
| p.L747_T751delinsP | 10 | 0.76 |
| p.E746_S752delinsA | 9 | 0.68 |
| p.E746_T751delinsI | 7 | 0.53 |
| p.E746_P753delinsVS | 6 | 0.46 |
| p.E746_T751delinsVP | 5 | 0.38 |
| p.T751_I759delinsN | 5 | 0.38 |
| p.E746_T751delinsIP | 4 | 0.30 |
| p.T751_E758del | 4 | 0.30 |
| p.E746_A750delinsAP | 3 | 0.23 |
| p.L747_A750del | 3 | 0.23 |
| p.L747_A750delinsS | 3 | 0.23 |
| p.E746_A750delinsP | 2 | 0.15 |
| p.E746_A750delinsQP | 2 | 0.15 |
| p.E746_E749del | 2 | 0.15 |
| p.E746_P753delinsLS | 2 | 0.15 |
| p.E746_T751del | 2 | 0.15 |
| p.E746_T751delinsV | 2 | 0.15 |
| p.E746_T751delinsVA | 2 | 0.15 |
| p.L747_A755delinsAT | 2 | 0.15 |
| p.L747_S752delinsQ | 2 | 0.15 |
| p.L747_T751delinsQ | 2 | 0.15 |
| p.T751del | 2 | 0.15 |
| p.A750_I759delinsPT | 1 | 0.08 |
| p.A750del | 1 | 0.08 |
| p.E746_A750delinsD | 1 | 0.08 |
| p.E746_A750delinsIP | 1 | 0.08 |
| p.E746_A750delinsRP | 1 | 0.08 |
| p.E746_E749delinsD | 1 | 0.08 |
| p.E746_E749delinsQ | 1 | 0.08 |
| p.E746_P753delinsIS | 1 | 0.08 |
| p.E746_S752delinsD | 1 | 0.08 |
| p.E746_S752delinsI | 1 | 0.08 |
| p.E746_T751delinsAPS | 1 | 0.08 |
| p.E746_T751delinsVS | 1 | 0.08 |
| p.K739_I744delinsN | 1 | 0.08 |
| p.L747_A750delinsEP | 1 | 0.08 |
| p.L747_A755delinsAN | 1 | 0.08 |
| p.L747_A755delinsSES | 1 | 0.08 |
| p.L747_A755delinsSKD | 1 | 0.08 |
| p.L747_A755delinsSMS | 1 | 0.08 |
| p.L747_E749delinsQ | 1 | 0.08 |
| p.L747_P753delinsQPS | 1 | 0.08 |
| p.L747_S752delinsQH | 1 | 0.08 |
| p.L747_S752delinsQP | 1 | 0.08 |
| p.L747_T751delinsN | 1 | 0.08 |
| p.L747_T751delinsPA | 1 | 0.08 |
| p.L747_T751delinsPR | 1 | 0.08 |
| p.L747_T751delinsS | 1 | 0.08 |
| p.P753_I759del | 1 | 0.08 |
| p.T751_A755del | 1 | 0.08 |
| p.T751_I759delinsS | 1 | 0.08 |
| Total | 1317 | N/A |

